# The Hidden Sweet Tooth of the Black Soldier Fly (*Hermetia illucens*)

**DOI:** 10.64898/2025.12.18.695165

**Authors:** Marie Merle, Tasnim Gueddes, Mouhamadou Moustapha Gueye, Meroua Foughar, Jessica Jiogue-Lacdo, Pierre-Olivier Maquart, Frédéric Marion-Poll, Jonathan Filée

## Abstract

The black soldier fly *Hermetia illucens* (BSF) is increasingly studied for its ability to convert organic waste into protein, offering solutions for waste valorisation and livestock feeding. While egg production depends on adult performance, the biology of adult BSF remains poorly understood. In particular, growing evidence suggests that adults can feed and benefit from sugar supplementation but the behavioural and molecular bases of sugar detection remain unknown.

To fill this gap, we tested whether adult BSF can detect and prefer sugar by using behavioural assays, binary-choice feeding tests and proboscis extension responses. We also described numerous gustatory sensilla on the proboscis with scanning electron microscopy, and characterized the responses to sugar in some of them with electrophysiological recordings. These experiments were complemented by an analysis of the gustatory receptor (GR) gene repertoire and the expression levels of sugar-responsive receptors.

All the experimental tests conducted in this study converge to show that the adults can detect and consume sucrose, with females responding more strongly than males. Genome analysis identified 28 GRs, a surprisingly small number for a generalist fly, including only three putative sugar-specific GRs homologous to those of the eight known sugar GRs in *Drosophila*. Moreover, one of these GRs show a general high level of expression including in the head and in the antennae whereas the two others display tissue-specific patterns of expression. We also identify a high number of GR pseudogenes, including four putative sugar receptor pseudogenes, indicating multiple gene loss events and substantial reduction of the GR repertoire compared to other dipteras sharing a similar ecological niche.

We conclude that although BSF possesses a reduced set of GR genes, including only three putative sugar-responsive GRs, adults have retained a strong sensitivity to sugars in their environment. The set of behavioural, morphological, and electrophysiological tools developed here provides a foundation for deeper investigations into feeding behaviour in this species of growing agroecological importance.

## Introduction

The Black Soldier Fly (BSF) (*Hermetia illucens*) has received considerable attention over the last decades due to its capacity to upcycle bio-waste into insect meal further used as feed for a large variety of animals such as poultry or fish (Wang and Shelomi, 2017). However, the emerging BSF industry is facing several challenges such as biosecurity concerns, market realities or welfare considerations (Lalander et al., 2025). To overcome these difficulties, there is a fundamental need for a better understanding of the BSF biology in order to improve BSF breeding.

Whereas the feeding behavior of the larva was extensively investigated, adult fly behavior remains poorly understood (Meneguz et al., 2023). For example, adult flies are considered as not feeding (Sheppard et al., 2002) but recent investigations have shown that adults have a functional digestive system and that food supplementation improves adults longevity (Bruno et al., 2019; Tettamanti et al., 2022). In particular, it was demonstrated that sugar supplementation in adult feeding affects various life history traits such as increasing egg production, extending the oviposition period or improving adult longevity (Barrett et al., 2025; Klüber et al., 2023; Macavei et al., 2020; Nakamura et al., 2016). The ability of adult BSF to benefit from sugar feeding may eventually imply that the fly has retained the capability to detect and taste carbohydrates in its natural environment. Accordingly, the mouthparts of these flies are covered with sensilla and adults display an extensible labellum (Bruno et al., 2019). These structures are known in other Diptera to be involved in chemoreception.

Strikingly, very few studies have addressed chemoreception in BSF. Whereas olfaction plays a crucial role in insect behavior, only two papers dealing with genetic analysis of chemosensory genes in BSF are available (Scieuzo et al., 2021; Xu et al., 2020). The process of chemosensory detection rely upon a number of gene families that include odorant-binding proteins (OBPs), chemosensory proteins (CSPs), sensory neuron membrane proteins (SNMPs), odorant receptors (ORs), ionotropic receptors (IRs) and gustatory receptors (GRs). Both studies on BSF are based on transcriptomic analyzes rather than complete genome analysis which can lead to under-estimation of the exact number of chemosensory genes. The study of Xu et al. identifies a limited set of 134 putative chemosensory genes including 27 OBPs (Xu et al., 2020). A transcriptomic analysis focused on odorant-binding proteins (OBPs) extended the OBP genes to 47 in the adult fly and they identified several volatile organic compounds that may interact with them (Scieuzo et al., 2021).

As for gustation, few behavioral studies showed that adult BSF preferentially chose to feed on sugar than water (Barrett et al., 2025; Romano et al., 2020). However, while the first genome of BSF provided reported 86 GRs (Zhan et al., 2020), only 10 GRs were found expressed (Xu et al., 2020). Such variation in the estimated numbers of GRs (10 to 86) is puzzling as this family of receptors plays a central role in coordinating insect feeding behaviors (van der Goes, van Naters and Carlson, 2006). By comparison, *Drosophila* genomes encode >60 GRs (Park and Kwon, 2011) and even highly food-specialized insects as some bees display a higher number of GRs (Fisher et al., 2025).

These limited data on BSF adult behavior and chemoreception prompted us to deploy original experiments to better understand sugar detection in this species. We combined binary-choice behaviour assay, proboscis extension response experiments, scanning microscopy and electrophysiological analyzes of labium sensilla to demonstrate that BSF are able to taste sugars. In addition, we provide whole-genome mining that identifies putative GR homologs in *Drosophila* that are known to mediate sugar tasting and show that these GR genes are highly expressed in the head of the flies. We conclude that although the BSF has retained a limited set of GR genes, and only three putative sugar-specific GRs, adult flies display a strong sensitivity to detect sugars in their environment.

## Material and Methods

### BSF Rearing

BSF adults were reared in a colony maintained in our laboratory. Larvae were reared on standard chicken feed adjusted to 50% moisture content under controlled environmental conditions (26 ± 1 °C, 60% relative humidity, 12:12 h light:dark photoperiod). The strain corresponds to the commercial strain used in the European insect industry (Guilliet et al., 2022) and is genetically very close to the reference genome assembly (Generalovic et al., 2021).

### Two-Choice Feeding Preference Assays

Drops of 50 mM sugar or water (2 µl) were deposited on a microscope glass slide and left to dry at 37°C overnight. The solution was added with 25% brilliant blue, which is a food dye (Brilliant blue FCF, Sigma Aldrich). Starved flies were introduced in a small cage enclosing three sugar and three water drops, placed over a light panel (white LEDs panel, Display Concept, Belgium). A webcam placed over the cages (Logitech HD920) was taking a picture at 20 s intervals using an open-source program (iSpy) running under Windows. We recorded the number of spots eaten over a 2 h period. The test was performed on recently emerged adult flies (less than 24 h). 25 females and 25 males were recorded.

### Proboscis Extension Response (PER)

Newly emerged adult BSF of both sexes starved during 24 h and immobilized in a tube, leaving the mouthparts exposed. Their proboscis was contacted with a toothpick dipped into a sugar solution for 1 s and we noted if the insect would extend its proboscis during 5 s. The stimulus was tested 3 times in a row, starting with the lowest sucrose concentration to the highest (from 10^-5^ M to 1 M). This test was performed on 25 females and 25 males.

### Scanning electron micrograph

BSF adults were euthanized by freezing at -20°C for 72 h. For the SEM observations, three males and three females were sacrificed. In some specimens the head or the proboscis was removed and mounted on aluminum stubs using double-sided carbon tape, critically-point dried and metalized with a thin layer of chromium and gold (8 nm). The cuticle morphology was analyzed using field emission scanning electron microscopy (SEM) with a Hitachi SU5000 microscope (MIMA2 platform, INRAE, Jouy-en-Josas, doi.org/10.15454/1.5572348210007727E12)

### Electrophysiological Assays

Adults were selected from the rearing room at approximately a day old. They were anesthetized on ice for several minutes and then disposed of on a magnetic stub using parafilm and strips of tape so that the proboscis was immobilized and held in an extended position. A drop of electrocardiogram gel was applied between the abdomen and a silver wire connected to the electrical ground, to serve as an indifferent electrode. After proper orientation of the preparation, selected individual taste sensilla were stimulated and recorded using the tip-recording method. A contact with the sensillum was maintained at least 2 s and consecutive stimulations to the same sensillum were separated by about 2 min to allow neurons to recover from the previous stimulation.

The recording electrode was a glass capillary connected to an amplifier with a silver wire, containing a solution with the stimulus and an electrolyte (30 mM tricholine citrate). The electric signal was recorded with a tasteProbe amplifier (DTP01, Syntech) (Marion-Poll and van der Pers, 1996), further amplified and filtered (10-2,000Hz) with a second amplifier (CyberAmp 320, Axon Instruments) and digitized at 10 kHz with a Data Translation card driven by a custom C++ program called dbWave2 (Marion-Poll, 1996). The spikes were detected and sorted using dbWave2, and the results were exported to a tabular file for further analysis.

### Whole-Genome Analysis

To identify gustatory receptor (GR) genes in BSF, we first assembled a database of GR protein sequences by performing a targeted literature survey across other dipteran species (supp Table 1). The Exonerate software (v.2.4.0) (Slater and Birney, 2005) was then used to align GR amino acid sequences from our database against the *H. illucens* genome (Generalovic et al., 2021; accession number GCF_905115235.1), using the protein2genome model. A maximum intron size of 3,000 bp was set for all GRs in order to maximize the detection of complete genes. Exonerate results were further analyzed with the InsectOR pipeline (Karpe et al., 2021), which detects chemoreceptors by identifying characteristic features such as the presence of seven transmembrane domains. Identified sequences were filtered using length criteria specific to GRs: only those between 800 and 1,600 nucleotides were retained and sequences identified as pseudogenes by InsectOR – due to frameshifts or premature stop codons – were discarded. The final curated sequences were combined with the initial database and subjected to multiple sequence alignment using Clustal Omega. The alignment obtained for GRs was used to build a maximum likelihood phylogenetic tree with IQ-TREE (Minh et al., 2020). The best-fitting substitution model was selected using ModelFinder (Kalyaanamoorthy et al., 2017) integrated in IQ-TREE. Branch support was evaluated using two complementary approaches: UFBoot with 1,000 replicates (Hoang et al., 2018) and the SH-aLRT test (Guindon et al., 2010).

### Three-Dimensional Structural Analysis

The three-dimensional structure of selected GRs was predicted using AlphaFold2 (Jumper et al., 2021), particularly for those orthologous to *D. melanogaster* receptors with experimentally characterized structures.

### Transcriptomic Analysis

Publicly available RNA-seq datasets of BSF were retrieved from the NCBI Sequence Read Archive (SRA). Larval transcriptomes were obtained from the Laval laboratory (accession numbers: SRR18283682, SRR18283691, SRR18283687), and adult transcriptomes were obtained from the Huazhong laboratory (accession numbers: SRR6656087, SRR6656088, SRR6656085, SRR6656086). Raw sequencing reads were processed with Kallisto v0.46.2 (Bray et al., 2016) for pseudoalignment and quantification of transcript abundances for each GR coding sequence. Quantification was carried out with default parameters. Transcript-level counts estimated by Kallisto were imported into edgeR v4.0.16 for downstream normalization and analysis. Raw counts were filtered to remove low-abundance transcripts prior to normalization. Expression levels were calculated as counts per million (CPM), scaled by library size. Expression profiles of GR transcripts across developmental stages and tissues were visualized as a heatmap. Heatmaps were generated using normalized log_2_(CPM + 1) values to highlight relative expression patterns of GRs across samples.

### Statistics

Data were analysed and plotted using LibreOffice (v.24.2.7) and R studio (v.2025.09.2). Statistical tests were performed using the Wilcoxon test.

## Results

### Analysis of BSF Sensitivity to Sucrose with Two-Choice Assays

To investigate how BSF flies selectively discriminate between a sucrose solution (100 mM) and water, we monitored the consumption of six blue droplets containing either water or a sucrose solution by a BSF enclosed in a small glass cage (Figure 1A). We estimated the number of droplets ingested over a one hour period by taking pictures every 20 s using a webcam (Figure 1B). Both male and female BSF demonstrated a strong preference toward sucrose compared to water. On average, each female consumed 2.4 droplets with sucrose per assay and males 1.6 (Wilcoxon test *P*-value between sexes = 0.037). Droplets made with water were systematically dismissed by both sexes leading to highly statistical differences between water and sucrose consumption for both sexes (Wilcoxon test *P*-value = 9.8e^-7^ and 2e^-10^ for males and females respectively). These findings suggest that adult BSF can detect and consume sucrose, and that this capacity seems more pronounced in females compared to males.

**Figure 1.**
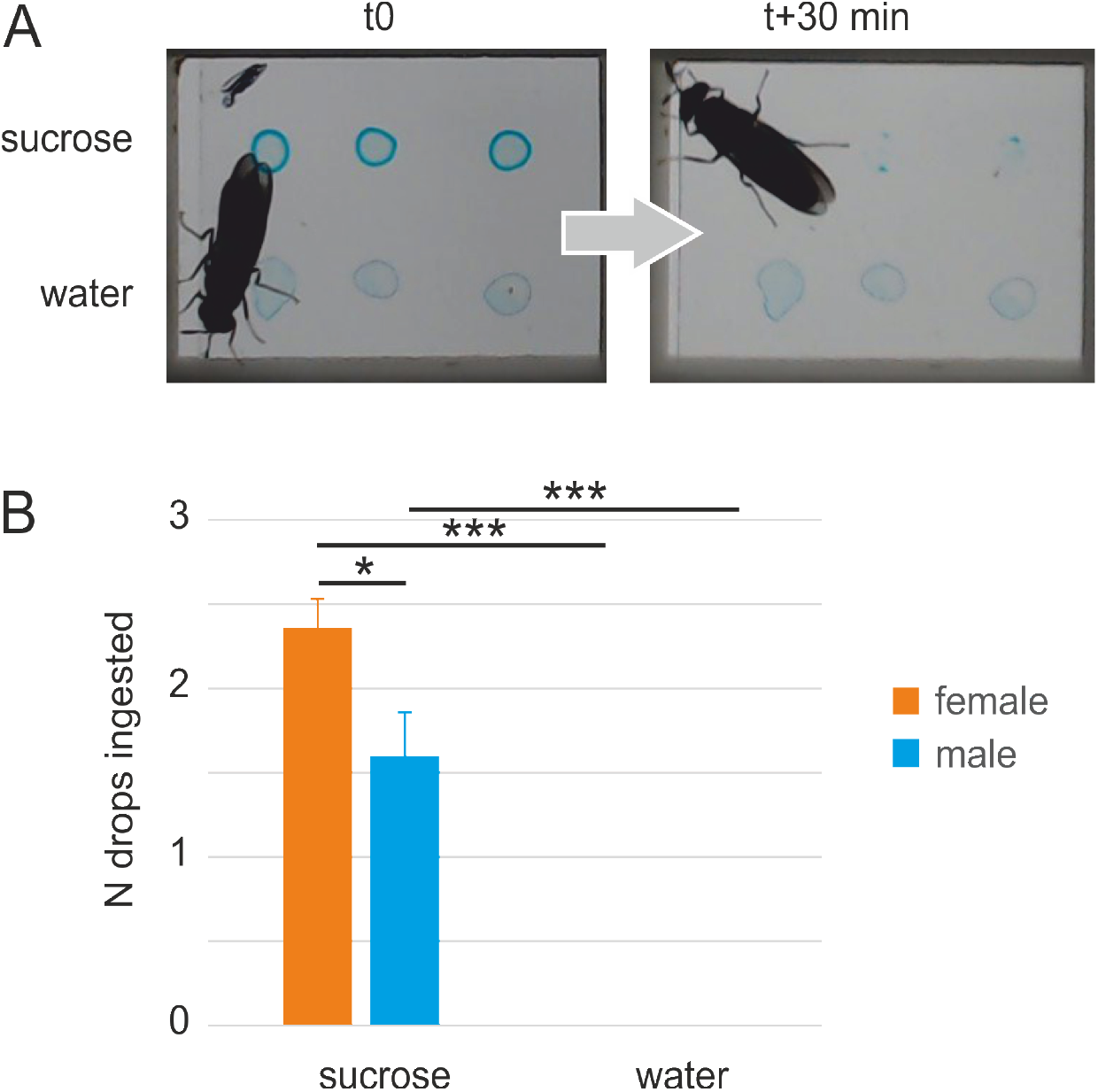
Two-choice feeding assay. Flies were given access to three drops of 100 mM sucrose or water mixed with brilliant blue dried on a microscope glass slide and the number of drops remaining after 1 h was recorded. (**A**) Representative sample of a fly given access to three drops of sugar and three drops of water at t0 and 30 minutes after. (**B**) Average count of drops consumed by flies after 1 h (mean +-s.e.m.; n = 25 individuals for each sex). Asterisks next to box plots indicate significant responses between sexes and between sucrose and water using Wilcoxon tests (\*\*\**P* < 0.001; \**P* < 0.05).

### BSF Gustatory Responses to Sucrose Using PER Assays

The proboscis extension reflex (PER) is an innate response in adult insects, in which they extend their mouthparts upon the application of a specific stimulus(Menzel and Müller, 1996). Here, PER assays were used to characterize the response of BSF males and females to sucrose solutions presented at different concentrations (Figure 2). This experiment shows that both males and females BSF extend their proboscis in a strong dose-dependent manner when their mouthparts are stimulated with a sucrose solution. Even low sucrose concentrations (1 mM) induce a behavioral response for both sexes. Indeed, the pattern of dose-response to sugar is very similar for both sexes, although males display a slightly lower response from 0.1 mM to 10 mM (differences between sexes for each sucrose concentration are not significant using a Wilcoxon test, *P* values > 0.05).

**Figure 2.**
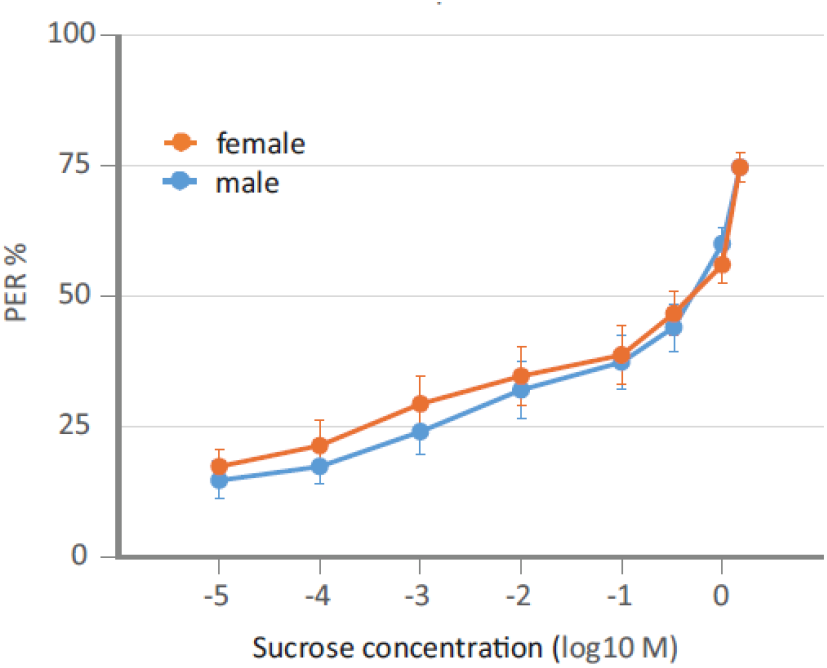
Proboscis extension reflex (PER) responses to increasing sucrose concentrations. Flies were immobilized into a holder and stimulated with a succession of increasing sucrose concentrations. Each point represents the percentage of responses obtained after three presentations of each stimulus in males and females (mean +-s.e.m.; n = 25 individuals of each sex).

### Scanning Electron Micrographs of the BSF Mouthparts

Scanning electron microscopy (SEM) observations of the BSF labium revealed a bilobed structure typical of dipteran mouthparts (Figure 3A). The external surface is densely covered with sensilla exhibiting heterogeneous morphologies, which can be divided into three main types : long (L), intermediate (I), and short (S) (Figure 3B). Their lengths range from approximately 50 µm for the shortest to 127 µm for the longest sensilla. Each sensillum has an apical terminal pore measuring up to 221 nm in diameter, suggesting a possible gustatory function (Figure 3C). The mean number of sensilla per labium was approximately 170 in males and 194 in females. The distribution of sensilla seems heterogeneous on the labium, with the L-type predominantly distributed in the distal region, whereas I and S-types were more frequent in the median portion of the labium.

**Figure 3.**
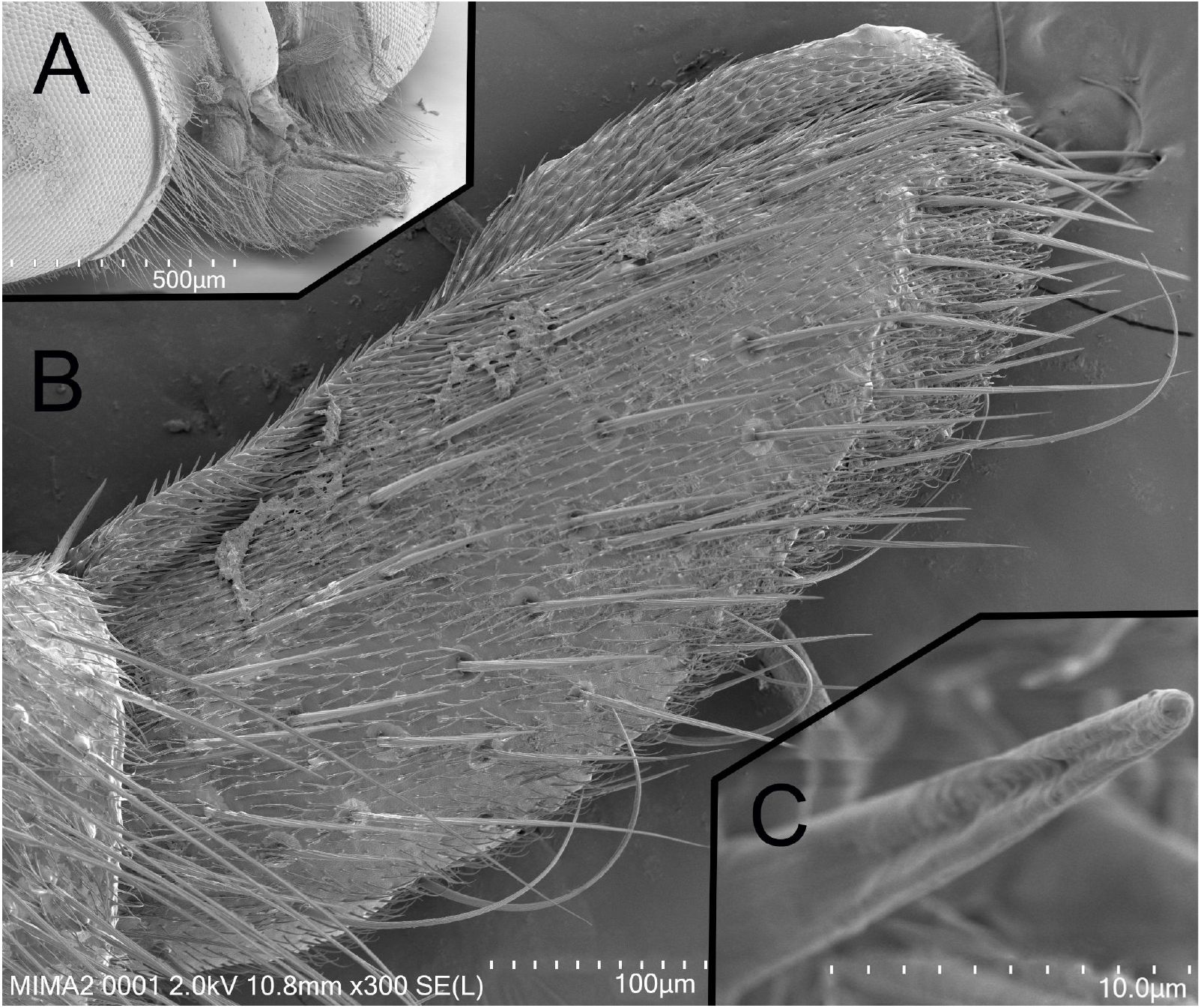
Scanning electron micrographs of adult BSF mouthparts. (**A**) Close-up view of the head (70×). (**B**) Detail of the right side of the labium (300×). (**C**) Tip of a gustatory seta showing the characteristic apical opening. Images were obtained with a scanning electron microscope (SEM) at the MIMA2 platform, INRAE.

### Electrophysiological Responses of L-type Chemosensilla to Sugars

We investigated the electrophysiological responses of the larger taste sensilla to increasing sucrose concentrations (Figure 4). L-type sensilla already showed significant responses to 1 mM sucrose with about 10 spikes per second (Figure 4A and 4B). Spike numbers sharply increase with sucrose concentrations for both sexes but females display a significant stronger response for intermediate sucrose concentrations (0.01 M and 0.1 M corresponding to 30 to 50 spikes per second in the females compared to 15 to 30 spikes per second for the males, Wilcoxon test *P* values = 0.026 and 0.019 respectively).

**Figure 4.**
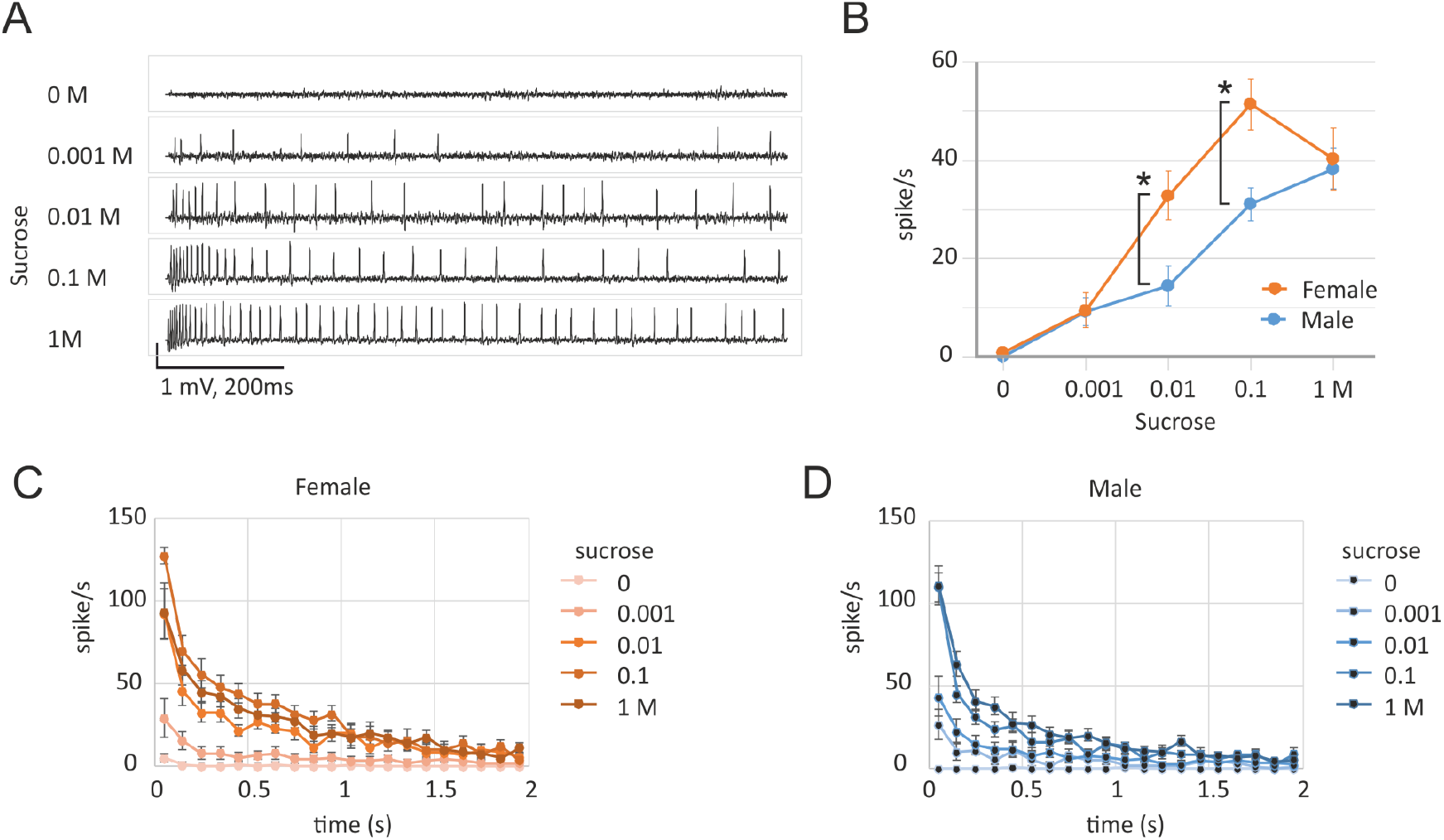
Electrophysiological responses to sucrose of individual L-type sensilla. **A**. Representative responses to increasing sucrose concentrations (with 30 mM tricholine citrate as an electrolyte). **B**. Average number of spikes recorded during 1 s. (mean +-s.e.m.) to increasing sucrose concentrations in females (orange circles; n = 10, 9, 7, 5, 8) and males (blue circles; n= 11, 9, 11, 8, 9). Asterisks next to the plots indicate significant responses between sexes (\**P* < 0.05;) using Wilcoxon test (two-sided). **C**. Time course of responses to increasing sucrose concentrations in females, obtained by counting the number of spikes in consecutive 100 ms bins (mean +-sem; n = 10, 9, 7, 5, 8). **D**. Time-course of the responses in males (mean +-sem; n = 11, 9, 11, 8, 9).

Moreover, the dose-response curve tends to reach a plateau at high sucrose concentration (1 M) for both males and females. Finally, the numbers of recorded spikes rapidly decreased across the time course of electrophysiological recordings, being halved at 0.5 s, and almost no recording is noticed after 1.5 s. The decreasing pattern is similar for males and females but, as noticed previously, females display higher potential changes compared to males for medium sucrose concentration.

In summary, all of our experimental tests converge to demonstrate that both BSF male and female display a strong dose-response tasting behavior to sucrose. This conclusion prompted us to analyze the putative genetic determinants of the sugar gustation in BSF by mining its genome to identify the putative gustatory receptor (GR) genes involved in this process.

### Gustatory Receptor Genes Identification and Functional Inference with other Dipteran GRs

A total of 28 GR genes were identified in the BSF genome (Supplementary Table 2). The phylogenetic tree revealed three predicted BSF GRs, grouped with nearly all sugar receptors of *D. melanogaster* (Figure 5, red). This suggests that these BSF GRs may share a similar function. Another clade, containing two predicted BSF GRs cluster within clades containing *D. melanogaster* GR63a and GR21a, respectively, both of which are involved in CO_2_ detection (Figure 5, green and orange clades). Likewise, two predicted BSF GRs clustered with major pheromone-related receptors (Figure 5, dark violet). In addition, one BSF GR was found to be phylogenetically close to *D. melanogaster* GR33a. Inspection of genomic coordinates (Supplementary Figure 1) revealed that GR genes of the same functional category are often physically clustered on the chromosome, and phylogenetically close. In some cases, clusters consist of multiple paralogs belonging to a single clade (*i*.*e*. HillGR13-15 sugar receptor clade in red, HillGR16-18 clade in blue, HillGR28-31 clade). This pattern indicates that many of these genes likely arose through tandem duplications, followed by divergence or pseudogenization.

**Figure 5.**
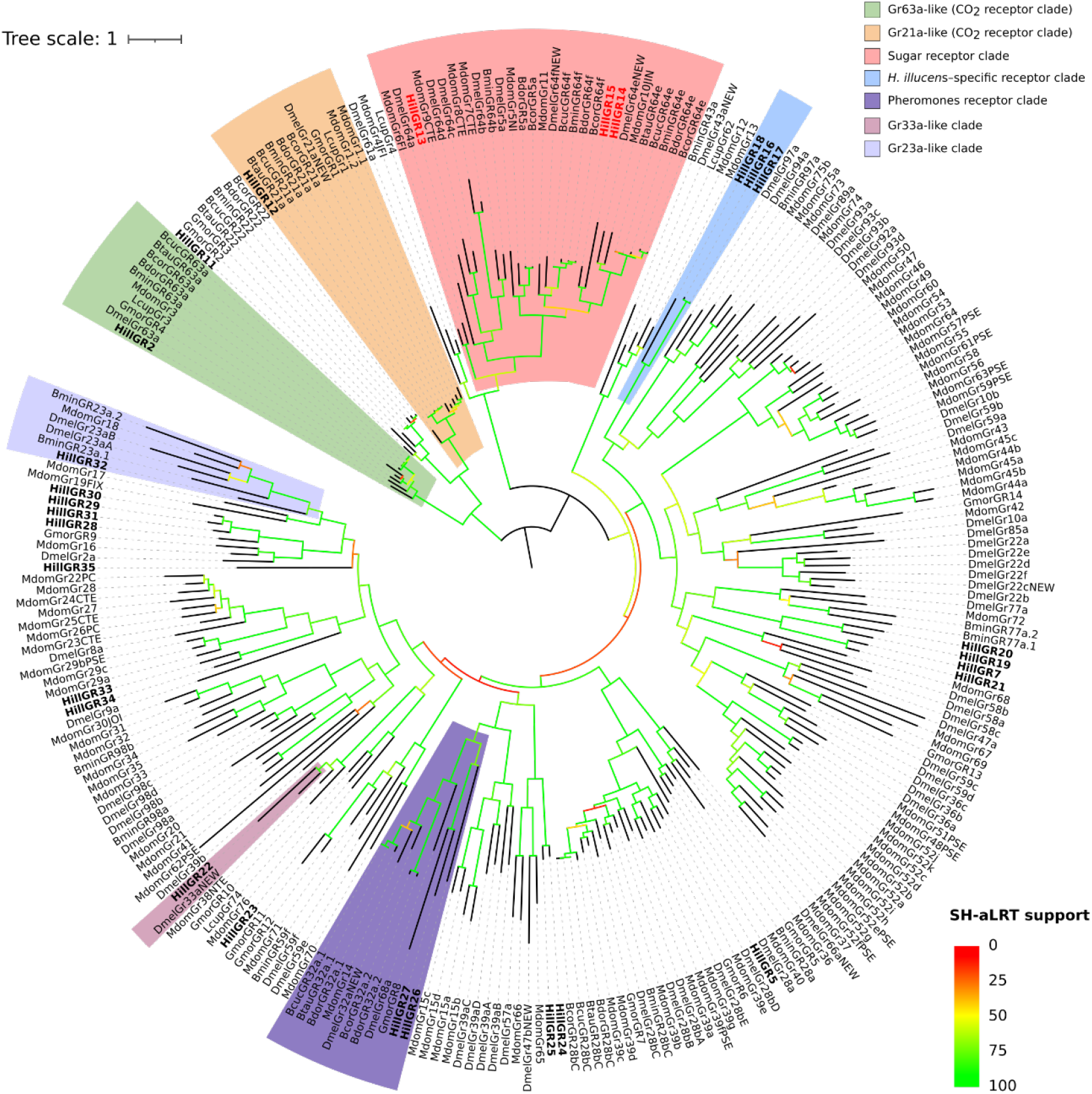
Maximum-likelihood phylogenetic tree of GR sequences from various dipteran species. The tree was constructed from 263 protein sequences (alignment length: 1,361 positions) using IQ-TREE. The substitution model selected for the analysis was VT+F+G4. By convention, the smallest clade containing the CO_2_ receptors GR63a and GR21a from *D. melanogaster* was used as the outgroup. Branch support values (SH-aLRT) are indicated by branch colors, ranging from red (lowest support) to green (highest support). Newly annotated BSF sequences are shown in bold. The clade of putative sugar receptors is enclosed in red, with BSF genes further highlighted in bold red labels.

Our GR identification pipeline also captured a number of GR genes with typical features of pseudogenes with the presence of frameshifts and smaller sizes compared to typical GR (Supplementary Figure 2). A subset of these GR pseudogenes, corresponding to 47 genes, displays an intermediate size between presumably intact GRs (coding DNA sequences > 800 bp) and small DNA remnants (CDS < 400 bp). Inclusion of these 47 GR pseudogenes in the phylogeny indicated that these pseudogenes generally fall close to putative functional BSF GRs (Supplementary Figure 3). For example, we identified a putative sugar GR pseudogene displaying a high level of relatedness with sugar BSF GRs HillGR14 and HillGR15. This observation is compatible with recent duplication events followed by a pseudogenisation. However, some of these pseudogenes seem to be distantly related from other BSF GR genes. Notably, we detected three GR pseudogenes located in the sugar GR clade that do not display any phylogenetic affinities with the three intact BSF HillGR13-15 sugar receptors. These GR genes might be the molecular relics of sugar receptors that were specifically lost in the BSF lineage.

### Detection of Sugar GRs in Different Developmental Stages and Tissues

The heatmap shows that about half of the GR genes are expressed (Figure 6). Among them, eight GRs are broadly expressed, being detected in almost all developmental stages and organs (at least 7 out of 8 conditions). Notably, larvae express a higher number of GRs overall than adults (14, 18, and 15 GRs across larval datasets versus 13, 14, 10, and 10 GRs in adults). Among the three putative sugar GRs (highlighted in red in the figure), one (HillGR15) is strongly expressed in all datasets. In contrast, HillGR13 and HillGR14 are detected only under specific conditions: HillGR13 in 20-day-old larvae and in one of the two whole-head datasets, and HillGR14 only in one of the two whole-head datasets. In adults, some GRs may be expressed outside the antennae, for example in the mouthparts (which are included in whole-head samples but where expression may appear diluted due to the high proportion of non-chemosensory transcripts), as well as in the legs or ovipositor. Mouthparts, antennae, and legs are all organs potentially involved in food detection.

**Figure 6.**
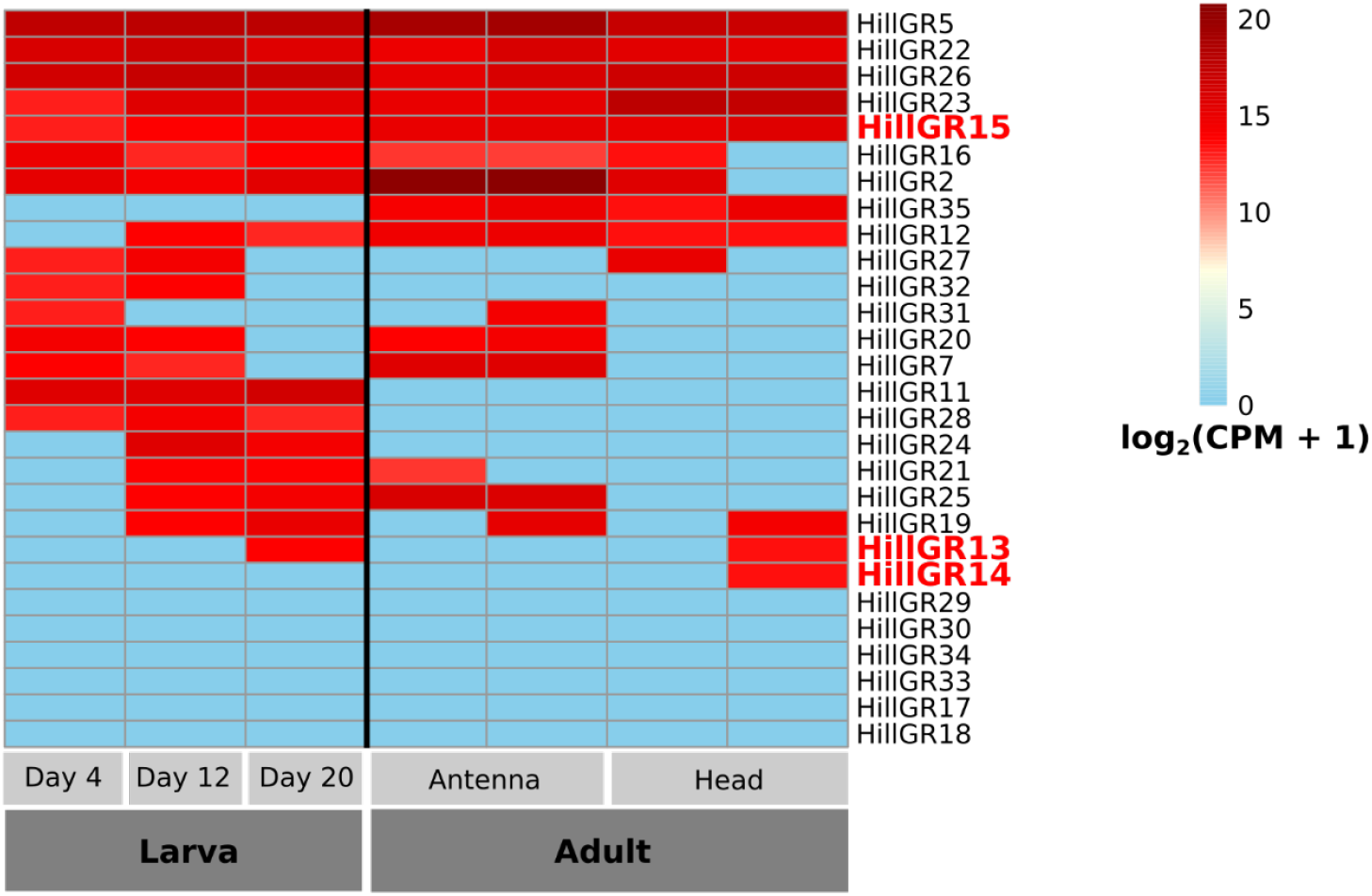
Heatmap of GR transcript expression across developmental stages and tissues in BSF. Expression values are shown as log_2_(CPM + 1), where CPM denotes counts per million normalized by library size. Color intensity reflects relative expression levels, from low (blue) to high (dark red). *Drosophila* homologs to sugar GRs found in BSF are highlighted in red.

### 2Structural Insights into GRs

Among *D. melanogaster* GRs, only GR43a and GR64a currently have available structural data obtained through crystallography (Ma et al., 2024). No clear ortholog of GR43a was identified in BSF. However, three predicted BSF GRs grouped with GR64a in the phylogenetic tree. One of these was successfully modeled with AlphaFold2, yielding a high-confidence structure (global pLDDT = 91.5; pTM-score = 0.905) (Figure 7A). Structural superposition with the recently resolved cryo-EM structure of *D. melanogaster* GR64a (PDB ID: 8JME, 2.5 Å resolution) resulted in a low RMSD of 1.067 Å (Figure 7B), indicating strong structural similarity and possible function conservation.

**Figure 7.**
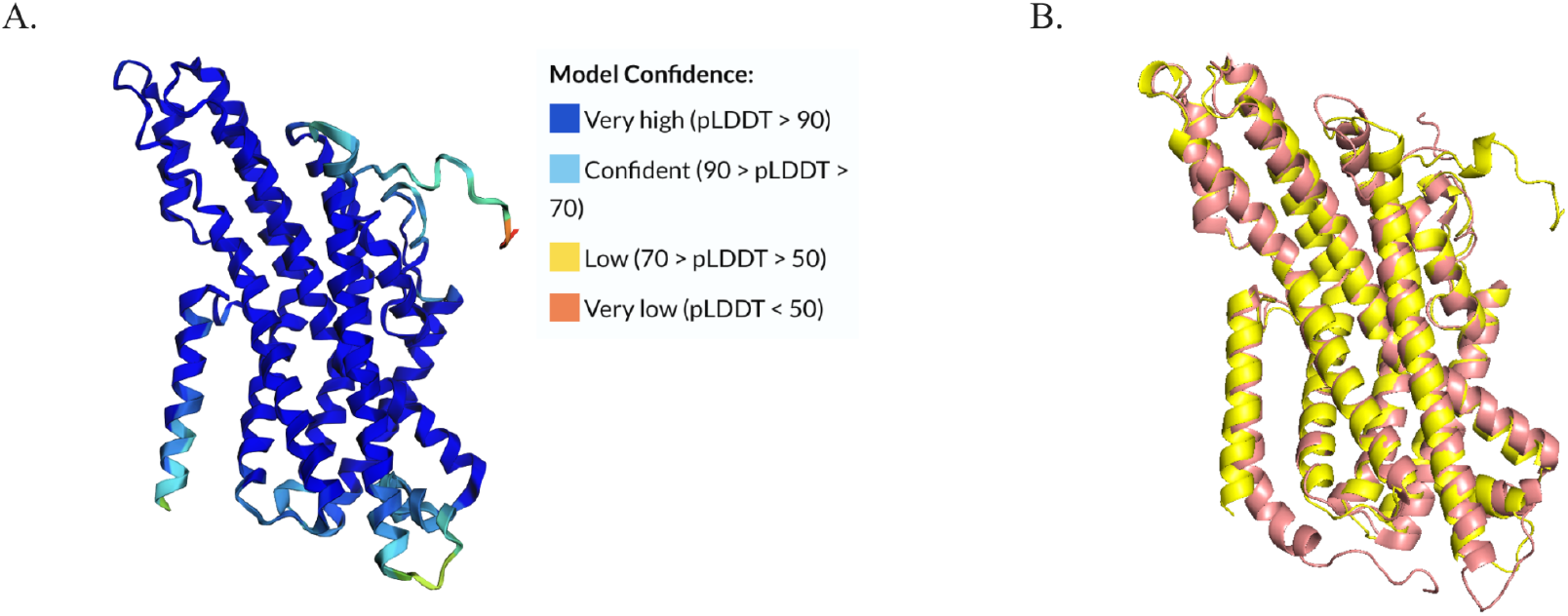
**(A)** Predicted 3D structure of the BSF GR HillGR13 (LR899010.1_120307204-120357024), generated using ColabFold v1.5.5 (based on AlphaFold2 with MMseqs2). Modeling parameters were set to num_relax = 0 and template_mode = none. Model quality metrics: mean pLDDT score = 91.5; predicted TM-score (pTM) = 0.905. **(B)** Superposition of the predicted HillGR13 structure with the *D. melanogaster* GR64a structure (PDB ID: 8JME; resolution: 2.5 Å; determined by cryo-EM, wild-type). The root mean square deviation (RMSD) between the two structures is 1.067 Å, indicating a high structural similarity.

## Discussion

The expanding industrial sector of edible insects for bioconverting waste into feed and food is now facing numerous challenges that mostly rely on a deficit of basic knowledge of the biology of the species. This is especially valid for the adult stage of most edible insects for which most of the corpus of knowledge focus on the larval stages. For example, it was commonly believed that adult BSF do not feed. But several lines of evidence support the idea that flies are able to drink and feed, to digest and benefit from the ingested food. Specifically, sugar supplementation of the water source influences some critical life-history traits such as extending the lifespan or increasing the fecundity of the flies suggesting that sugar consumption is beneficial (Barrett et al., 2025; Klüber et al., 2023; Macavei et al., 2020; Nakamura et al., 2016). However if flies can benefit from sugars, they must first detect and taste them in their environment. This was the starting point of our study with the goal to evaluate the functional capacity of the BSF to taste sucrose and if so, to gain insight into the genetic architecture of sugar sensing.

### Adults BSF Love Sugar

We first combined two-choice behavioural tests and proboscis extension reflex (PER) assays to evaluate the capacity of adult BSF to taste sugars. Our observations indicate that taste sensilla on the proboscis exhibit strong responses to sucrose, even at 1 mM. Both sexes respond positively when stimulated with sugars but females seem more prone to taste sucrose. Morphological observations of the BSF labium using scanning electron microscopy revealed a high abundance of gustatory sensilla, suggesting a central role of this structure in detecting sapid stimuli. These findings confirm the presence of a functional gustatory system in adults, supporting their ability to detect and discriminate among chemical cues and contradicting earlier assumptions that gustatory capacity is reduced in the adult stage. The mean number of sensilla per labium was notably high (170 in males, 194 in females) compared to *D. melanogaster* (31; Weiss et al., 2011). Compared with this insect model, the BSF exhibits both a higher density and greater morphological diversity of labial sensilla, suggesting its refined adaptation to gustatory perception in its environment.

The high density of long-type sensilla on the fly’s labium prompted us to study their electrophysiological responses when exposed to a sucrose solution. In line with the previous results, dose–response curves showed that the threshold sucrose concentration is low for the long type labium sensilla (as of 1 mM). Sucrose stimulation induced a strong response for both sexes, but as noticed with two-choice and PER assays, females displayed a slightly stronger response for medium sucrose concentration (10 to 100 mM). It is tempting to propose that females are more prone to detect sugar in relation to the energy investment required for egg production compared to male. Although female BSFs are larger than males, they live shorter lives, although nutrients gathered from males during mating increase their lifespan. (Harjoko et al., 2023).

Similarly, adding sugar to the water increased male lifespan threefold, but female lifespan only twofold, supporting higher nutrient requirements in females (Nakamura et al., 2016). Our data demonstrate that male and female BSF are able to taste sucrose, even if the adult flies are not known to actively seek food in nature as observed in some other insects. Interestingly, even hematophagous insects considered specialized still feed on sugars when their main food source is not available. For instance, mosquitoes (Yuval, 1992) and triatomines (Da Lage et al., 2024) consume plants to survive periods without hosts.

### Reduced GR Repertoire in BSF

In addition to these experimental tests, we have mined the BSF reference genome to identify the possible molecular determinants of sugar tasting. We identified 28 putative GRs which is significantly higher than the 10 GRs identified on transcriptomes in a previous study (Xu et al., 2020). But this is much lower than the 86 GRs in the study of Zhan *et al*. (Zhan et al., 2020).Our bioinformatic GR identification pipeline has identified numerous large GR pseudogenes (47 genes) in the BSF genome that could explain this difference. Unfortunately, the publication of Zhan *et al*. lacks any supporting data, including the 86 GR sequences they have identified making it impossible to compare and understand the origin of the two different estimations.

By comparison with other insects, the GR gene repertoire of the BSF appears highly reduced. For instance in other diptera such as *D. melanogaster* or the house fly *Musca domestica*, much more GR genes are encoded (68 and 79 respectively). Reduction of chemoreceptor repertoire has been associated with habitat specialisation. For example GRs repertoire reduction in links with adaptations for underground life in crickets (Chudhary et al., 2025). Food specialisation has also generated a similar pattern. For example, the nectar specialization observed in some solitary bees has led to a highly reduced GR gene repertoire, with as few as 10 GRs encoded in the solitary bee *Cerceris arenaria* genome (Obiero et al., 2021). Some strictly hematophagous insects such as the tsetse fly also encode a small GR repertoire (17 genes; Obiero et al., 2014).

As BSF are known to exploit an astonishing diversity of plant and animal food sources – which would theoretically require more receptors to evaluate – one would expect a more diversified GR repertoire than observed. It could also be selectively advantageous for an invasive species to be able to detect diverse food sources by encoding a large number of GR genes as observed in invasive generalist ant species, for instance (Smith et al., 2023). But again, it’s paradoxical that a cosmopolitan species such as the BSF, that is known to have invaded the world in the last centuries, encodes such a limited set of GRs. We can propose some hypotheses to explain the BSF GR repertoire reduction. First, the BSF ecology in its native range in America is poorly known, but it sounds plausible that BSF have been adapted to specific food sources which may have led to a GR repertoire reduction. Alternatively, the domesticated BSF strain used for genome sequencing may have undergone some events of GR gene loss in links with the domestication process. Indeed, domestication corresponds to a kind of a food specialisation and niche adaptation in which many functions required in the wild become unnecessary in captive environments. For instance, domestication of cows and horses is associated with events of pseudogenization of multiple bitter taste receptor genes (Dong et al., 2009). This explanation is consistent with the high number of GR pseudogenes identified in the BSF genome but further confirmation is needed by analyzing other BSF genomes, including those from wild specimens.

### Evolution of Sugar Receptors Recapitulates Reduction of the Entire GR Repertoire

Among this reduced set of 28 GRs, three GRs are phylogenetically close to the sugar receptors of *D. melanogaster* (HillGR13, HillGR14 and HillGR15). By comparison, *D. melanogaster* and *M. domestica*, which occupy a similar anthropo-ecological niche, display much more sugar receptor genes (9 and 7 respectively). HillGR14 and HillGR15 are closely related to the GR64e gene in *D. melanogaster*, which is required to detect fermented products as glycerol (Wisotsky et al., 2011). HillGR13 is phylogenetically close to the *D. melanogaster*’s sugar receptors GR64b/c/d that both contribute to sugar detection (Fujii et al., 2015). Four putative sugar GR pseudogenes have also been detected in the BSF genome. Three of them do not display close phylogenetic affinities with the BSF intact sugar receptors (HillGR13-15). The small sugar GR repertoire in BSF, along with several sugar GR pseudogenes, may reflect a more general process of GR gene loss and repertoire contraction in the BSF genome.

In addition, HillGR13 displays strong predicted three-dimensional structural similarity with the sugar receptor GR64a. While phylogenetic analysis is primarily based on DNA or protein sequence comparisons, protein tertiary structure is often more conserved than primary sequence. More than 50% of amino acids can vary without significantly altering overall protein structure (Schaefer and Rost, 2012). Comparative studies of secondary structure, residue contacts, and solvent accessibility have shown that structural features are 3-10% more conserved than sequences (Illergård et al., 2009). Thus, structural analyses can provide complementary insights into receptor function, particularly in multigene families.

It should be noted that HillGR14 and HillGR15 appear as sister taxa in the tree and they are located as a tandem in the chromosome 2, suggesting a recent clade-specific gene duplication. Whereas HillGR15 gene is highly expressed in most larval and adult tissues, HillGR14 displays a narrower gene expression pattern in the head of the adult. This observation is compatible with a gene duplication event followed by a regionalization of the expression of one of the two paralogs in some tissues or in some developmental stages, whereas the other copy remains transcribed constitutively. As observed in *Drosophila*, the presence of multiple sugar GRs may also be explained by a degree of specialization for specific types of sugar (Fujii et al., 2015a).

For example, GR64a in *Drosophila*, which displays high structural similarities with HillGR13, is specifically activated by disaccharides (as sucrose and maltose) (Ma et al., 2024). A combination of several GRs is also sometimes required to taste specific sugars (Slone et al., 2007). Here, we have tested only sucrose and it should be valuable to test other kinds of saccharide to evaluate the exact BSF sugar sensibility. Ultimately, the generation of defective mutants by genome editing of the three BSF sugar GRs will validate the exact functional roles of each of these receptors.

Finally, the apparent reduction of sugar GRs in BSF can be further contextualized by comparative genomic analyses of gene family expansions and structural variants. Studies have shown that BSF exhibits specific gene duplications in families related to olfactory and immune functions, while other Stratiomyidae show expansions in digestive and metabolic genes, reflecting their ecological roles as decomposers (Zhou et al., 2025). This pattern suggests that rather than a general expansion of all chemoreceptors, BSF may have retained a limited set of sugar GRs while maintaining duplications in other gene families that optimize detection of relevant cues and processing of diverse organic substrates. In this context, the reduced sugar GR repertoire does not necessarily imply a limitation in ecological adaptability but may represent a functional specialization shaped by selective pressures on efficient resource exploitation.

## Conclusion

Although the BSF genome encodes a limited set of three different putative sugar GRs, our experimental setup provides compelling and original evidence that the adult BSF are indeed able to taste sugar. The high abundance of GR pseudogenes in the BSF genome is compatible with a reduction of the global GR repertoire, including a substantial sugar GR gene contraction process compared to other dipteras. Future prospects on the exact function of these three putative sugar GRs would be valuable to understand the evolution of BSF sugar detection.

Moreover, putative GRs involved in the detection of repulsive chemicals and pheromones that regulate male courtship in *D. melanogaster* have been identified in the BSF genome (GR33a and GR32a/GR68a) (Miyamoto and Amrein, 2008; Moon et al., 2009). Similarly, genes encoding CO_2_ receptors in *D. melanogaster* have closely related homologs in the BSF genome (Kwon et al., 2007). We believe that the behavioral and electrophysiological experimental design developed in this study may be suitable for testing a wide variety of chemical substances, with the goal of gaining further insights into BSF chemoreception.

## Abbreviations

BSF: black soldier fly
CDS: coding DNA sequence
CO_2_: carbon dioxide
GR: gustatory receptor
RNA-Seq: RNA sequencing
SEM: scanning electron microscopy
SH-aLRT: Shimodaira–Hasegawa-like approximate likelihood ratio test.

## Acknowledgments

We would like to thank all the crew involved in BSF rearing at EGCE lab (Rémi Jeannette, Chloé Genevey, Philene Corinne Aude Um Nyobe and Mhant Kondé), Céline Moreno and Béatrice Denis for their help during the behavioral tests. We would like to thank Dr. Vlad Costache and the MIMA2 Imaging Core Facility, Microscopie et Imagerie des Microorganismes, Animaux et Aliments, INRAE, Jouy-en-Josas for their help around the SEM.

## Authors Contribution

JF designed and directed the project; MM conceived and carried out the bioinformatics analyzes with the help of MMG; FMP and POM conceived the experimental setup; TG, JL, MF, FMP, POM and JF performed the experimental tests and analyzed the data; MM, POM, FMP and JF wrote the article.

## Data and Code Availability

Raw data are available on Zenodo at https://zenodo.org/communities/anr_insection/, under the DOIs 10.5281/zenodo.17881277 (two-choice feeding assay records, PER records, electrophysiological records and R codes for statistical analysis), 10.5281/zenodo.17899547 (for SEM raw images) and 10.5281/zenodo.17901058 (for sequences, tree files, quantification of GR expression, and gene name correspondence).

