## Supplementary Figures and Tables for "The Hidden Sweet Tooth of the Black Soldier Fly (*Hermetia illucens*)"

### Supplementary Materials

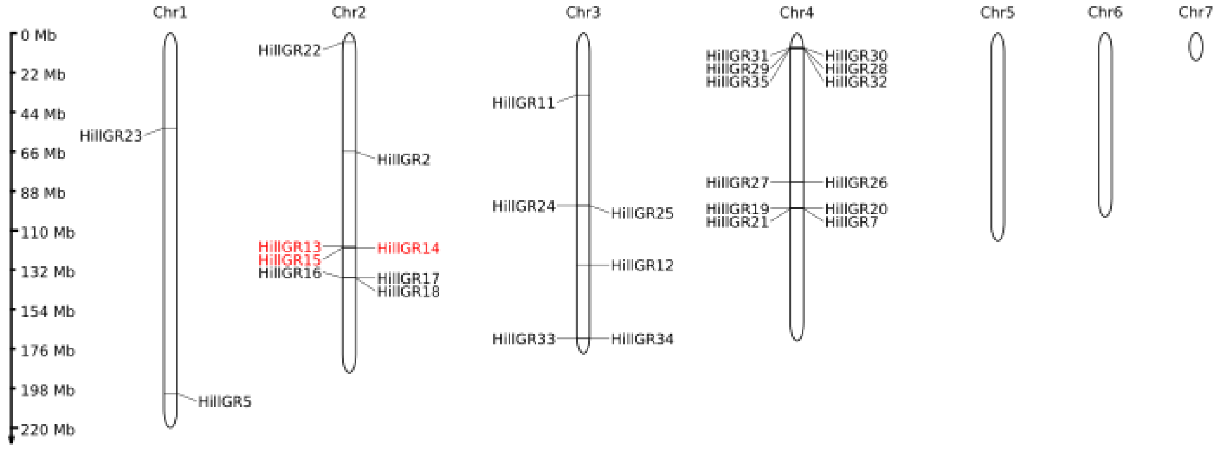


**Supplementary Figure 1. Chromosomal localization of BSF GR genes**. Each chromosome is represented as a vertical bar, with gene positions indicated along the length of the chromosome. GRs corresponding to sugar receptors are colored in red. The figure provides an overview of the genomic distribution of GRs, allowing visualization of potential clusters and relative positions across the genome.

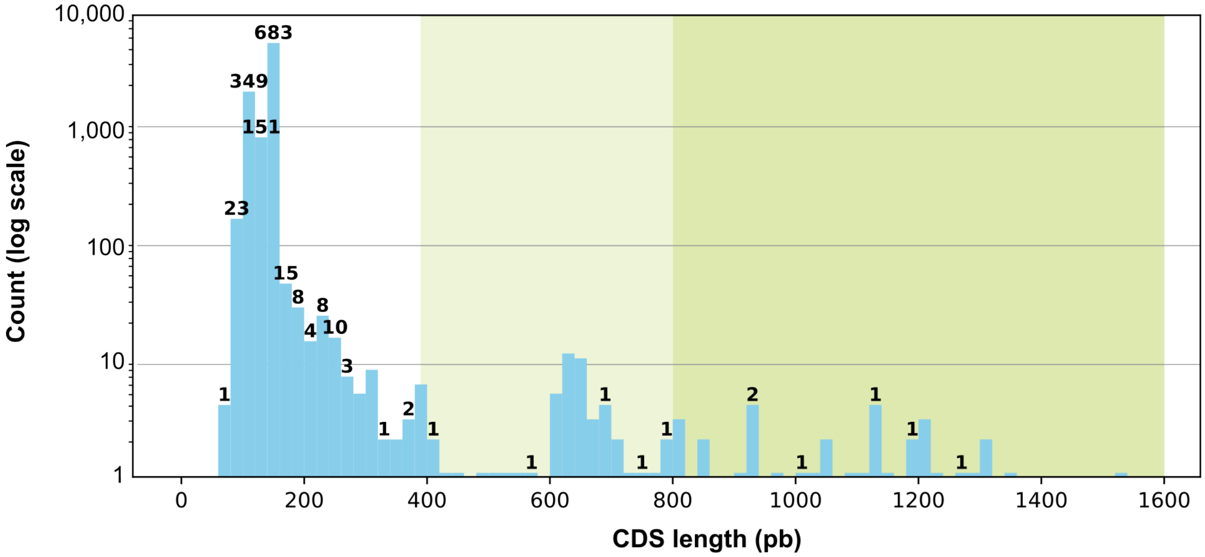


**Supplementary Figure 2. Length distribution of BSF CDSs predicted by the InsectOR pipeline before size filtering and manual curation.** The histogram shows the number of CDSs per length class (Y-axis on a logarithmic scale). Numbers above the bars indicate sequences flagged as pseudogenes by InsectOR. The dark-green portion represents CDSs retained in the final annotation (excluding pseudogenes), while the combined light- and dark-green portions correspond to sequences used for the complementary phylogenetic analysis (shown in Supplementary Figure 3). The white portions indicate CDSs discarded for being too short.

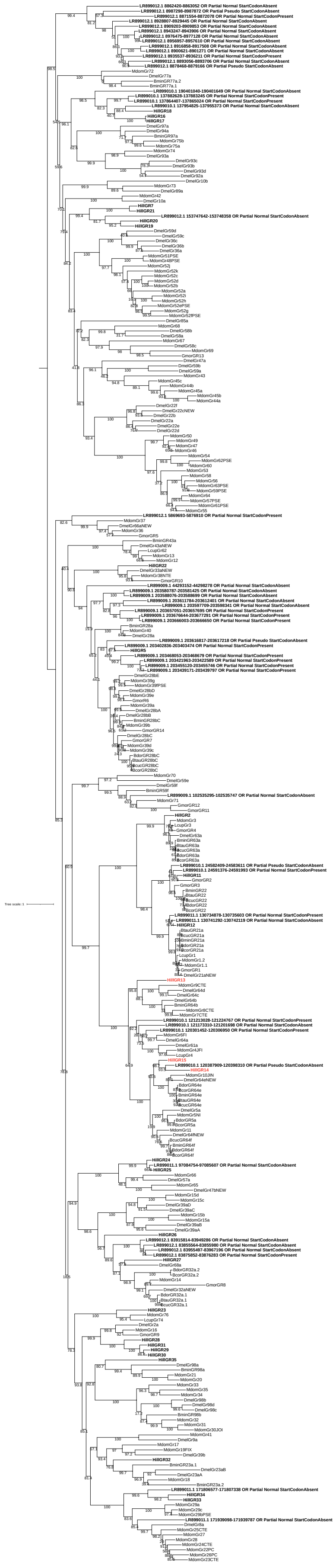
Tree scale: 1

**Supplementary Figure 3. Maximum-likelihood phylogenetic tree of GR sequences from various dipteran species, including short amino acid sequences (from 130 aa) and translated CDS classified as pseudogenes in the InsectOR output.** The tree was inferred from 312 protein sequences (alignment length: 1,352 positions) using IQ-TREE under the JTT+F+R8 substitution model and midpoint-rooted. Newly annotated *H. illucens* sequences are highlighted in bold. The putative *H. illucens* sugar receptors are shown in red.

**Supplementary Table 1. GR in selected dipteran species.** This table lists the number of GRs identified in various dipteran species. For each species, the taxonomic family, the number of GRs, the type of molecular data available (genome or transcriptome), and the corresponding literature reference are provided. These sequences were collected to serve as a reference for identifying GRs in the BSF genome.

| **Species** | **Family** | **GR** | **Data Type** | **Reference** |
|---|---|---|---|---|
| *Bactrocera correcta* | Brachycera; Muscomorpha | 8 | Transcriptome | Wu et al., 2020 |
| *Bactrocera dorsalis* | Brachycera; Muscomorpha | 7 | Transcriptome | Wu et al., 2020 |
| *Bactrocera minax* | Brachycera; Muscomorpha | 17 | Transcriptome | Wu et al., 2020 |
| *Drosophila melanogaster* | Brachycera; Drosophilidae | 60 | Genome | Robertson et al., 2003; Hekmat-Scafe et al., 2002 |
| *Glossina moristans* | Brachycera; Muscomorpha | 14 | Genome & Transcriptome | Obiero et al., 2014 |
| *Lucilia cuprina* | Brachycera; Muscomorpha | 5 | Transcriptome | Wulff et al., 2014 |
| *Musca domestica* | Brachycera; Muscomorpha | 79 | Genome | Scott et al., 2014 |
| *Zeugodacus cucurbitae* | Brachycera; Muscomorpha | 7 | Transcriptome | Wu et al., 2020 |
| *Zeugodacus tau* | Brachycera; Muscomorpha | 6 | Transcriptome | Wu et al., 2020 |

**Supplementary Table 2. Annotation of GR genes.** This table provides the genomic coordinates of each GR gene annotated in this study.

| Gene name | Start (bp) | End (bp) | Chromosome | Contig/Accession |
|---|---|---|---|---|
| HillGR2 | 66,403,317 | 66,406,227 | 2 | NC_051850.1 |
| HillGR5 | 203,380,985 | 203,391,551 | 1 | NC_051849.1 |
| HillGR7 | 98,452,864 | 98,457,453 | 4 | NC_051852.1 |
| HillGR11 | 35,121,710 | 35,123,214 | 3 | NC_051851.1 |
| HillGR12 | 130,510,219 | 130,511,806 | 3 | NC_051851.1 |
| HillGR13 | 120,307,204 | 120,357,024 | 2 | NC_051850.1 |
| HillGR14 | 120,414,420 | 120,420,796 | 2 | NC_051850.1 |
| HillGR15 | 120,435,886 | 120,461,073 | 2 | NC_051850.1 |
| HillGR16 | 137,920,339 | 137,929,188 | 2 | NC_051850.1 |
| HillGR17 | 137,909,714 | 137,910,691 | 2 | NC_051850.1 |
| HillGR18 | 137,938,433 | 137,939,620 | 2 | NC_051850.1 |
| HillGR19 | 98,418,680 | 98,430,105 | 4 | NC_051852.1 |
| HillGR20 | 98,436,193 | 98,437,107 | 4 | NC_051852.1 |
| HillGR21 | 98,442,643 | 98,443,671 | 4 | NC_051852.1 |
| HillGR22 | 5,322,467 | 5,328,364 | 2 | NC_051850.1 |
| HillGR23 | 53,439,288 | 53,446,866 | 1 | NC_051849.1 |
| HillGR25 | 97,108,785 | 97,109,785 | 3 | NC_051851.1 |
| HillGR24 | 97,155,999 | 97,156,968 | 3 | NC_051851.1 |
| HillGR26 | 84,004,124 | 84,010,274 | 4 | NC_051852.1 |
| HillGR27 | 83,684,356 | 83,685,276 | 4 | NC_051852.1 |
| HillGR28 | 8,100,437 | 8,101,850 | 4 | NC_051852.1 |
| HillGR29 | 8,089,079 | 8,090,151 | 4 | NC_051852.1 |
| HillGR30 | 8,072,198 | 8,073,655 | 4 | NC_051852.1 |
| HillGR31 | 8,066,115 | 8,067,536 | 4 | NC_051852.1 |
| HillGR32 | 8,146,831 | 8,148,195 | 4 | NC_051852.1 |
| HillGR33 | 171,771,102 | 171,771,917 | 3 | NC_051851.1 |
| HillGR34 | 171,790,009 | 171,790,821 | 3 | NC_051851.1 |
| HillGR35 | 8,111,048 | 8,116,264 | 4 | NC_051852.1 |
